# An Indian Diet–Relevant Rat Screening Model for Hypertriglyceridemia-Associated Fatty Liver

**DOI:** 10.64898/2026.02.15.705955

**Authors:** K Saranya, Priyanka Jadhav, Sadiya Mehaboob, Shreya Shahapur, Gopi Kadiyala, Uday Saxena

**Author notes:** Corresponding author : Uday Saxena.

## Abstract

Hypertriglyceridemia is a dominant and early metabolic abnormality underlying fatty liver disease in Indian populations, often preceding obesity, insulin resistance, or inflammatory liver injury. Many diet-induced rodent models of hepatic steatosis rely on extreme obesogenic or fructose-rich diets that poorly reflect real-world Indian dietary patterns. Here, we describe a diet-induced rat screening model designed to reflect typical Indian cereal-rich, visible-fat dietary exposure and to preferentially induce triglyceride-centric hepatic lipid accumulation. The model reproducibly induces hepatic triglyceride deposition with preserved liver architecture and minimal inflammatory features, aligning with early-stage fatty liver observed clinically in Indian patients. This work does not propose a novel disease model nor evaluate therapeutic efficacy, but establishes a translationally relevant screening tool for prioritizing lipid-modulating interventions in hypertriglyceridemia-associated fatty liver.

We show that the high-fat diet increased serum triglycerides ∼1.8 -fold versus chow (normalized index 1.0 vs 1.8), with organ weights remaining within ∼0.95–1.00 of reference (normalized indices), supporting screening tolerability. Secondary changes in liver morphology and histopathology were indicative of fatty liver.

## Introduction

Fatty liver disease has emerged as a major public health challenge in India, driven by urbanization, dietary transition, and a rising burden of dyslipidemia. Indian patients frequently develop hepatic steatosis at lower body mass indices than Western counterparts, with hypertriglyceridemia playing a central pathogenic role. Conventional rodent models often emphasize obesity, insulin resistance, or advanced inflammatory disease, limiting their relevance for early-stage, diet-driven fatty liver common in Indian populations. There is therefore a need for screening models that reflect Indian dietary patterns and triglyceride-centric hepatic lipid accumulation.

## Materials and Methods

### Animal Model, Husbandry, and Diet

All animal procedures were approved by the Institutional Animal Ethics Committee (IAEC) under CPCSEA guidelines. Specific pathogen–free male Wistar rats (7–8 weeks of age; body weight 170 ± 20 g at randomization) were used in this study. Animals were procured from Hylasco Biotechnology India Pvt. Ltd. (CPCSEA Registration No. 1808/PO/RcBt/S/15/CPCSEA).

Animals were housed individually in stainless steel cages under controlled environmental conditions (temperature 22 ± 3 °C; relative humidity 30–70%; 12 h light/12 h dark cycle) with a minimum 7-day acclimatization period prior to randomization. Purified Aqua-guard drinking water was provided ad libitum.

To induce hypertriglyceridemia-associated fatty liver reflective of Indian dietary patterns, rats were fed ad libitum a cereal-rich, saturated fat–enriched high-fat diet containing cholesterol, sodium cholate, lard, and sucrose for 8 consecutive weeks. This diet reproducibly induces triglyceride-centric hepatic steatosis with preserved liver architecture and minimal inflammation, consistent with early-stage fatty liver disease in Indian populations. Control animals, where applicable, received standard pelleted rodent chow.

The dietary screening model resulted in elevated serum triglycerides, consistent hepatic lipid accumulation predominantly in the form of macrovesicular steatosis, minimal inflammatory changes, and organ weights within physiological ranges, supporting its suitability for screening and intervention studies.

### Animals

#### Diet Composition and Induction Strategy

The control group received a standard laboratory chow diet supplied. The experimental group was fed a lipid-enriched diet formulated to induce hypertriglyceridemia-associated hepatic steatosis. The high-fat diet consisted of cholesterol (2.0%), sodium cholate (0.5%), lard (10.0%), propylthiouracil (0.2%), and sucrose (5.0%), with the remaining 82.3% composed of a basal feed.

The basal feed comprised corn flour (36%), wheat flour (35%), wheat bran (15%), soybean flour (10%), yeast powder (1%), sodium chloride (1%), bone powder (1%), and fish liver oil (1%). This cereal-rich, saturated fat–enriched formulation reflects key elements of contemporary Indian dietary patterns, including reliance on wheat-based staples combined with visible fats analogous to ghee and butter, and increasing refined sugar intake. The diet was selected to create a metabolic environment characterized by chronic triglyceride overload rather than extreme caloric excess or advanced inflammatory stress.

#### Metabolic and Histological Assessment

Animals were monitored for general health and body condition throughout the feeding period. Blood samples were collected under standardized conditions for assessment of circulating triglycerides using validated enzymatic assays. At study termination, livers were excised, weighed, and processed for hepatic triglyceride quantification and histological analysis. Liver sections were stained with hematoxylin and eosin, and lipid-specific staining with oil red O was performed where appropriate.

## Results

**Table 1.** Comparison of the Rat Screening Diet with Typical Indian Dietary Patterns.

| Dietary Component | Rat Screening Diet | Typical Indian Diet (Urban/Semi-Urban) |
| --- | --- | --- |
| Staple carbohydrates | Corn flour, wheat flour, wheat bran | Wheat (chapati), rice, millets |
| Primary fat source | Lard (10%) | Ghee, butter, vanaspati, refined vegetable oils |
| Sugar content | Sucrose (5%) | Refined sugar, sweets, sweetened beverages |
| Protein sources | Soybean flour, yeast powder | Pulses, legumes, dairy |
| Micronutrient lipids | Fish liver oil | Animal fats, dairy fats |
| Dietary pattern emphasis | Triglyceride-centric lipid overload | Visible fat intake with cereal-rich base |

Shown below in Table 2 are the key findings. The dietary regimen reproducibly induced serum and hepatic triglyceride accumulation across animals. Histological examination revealed predominantly macrovesicular steatosis with preserved lobular architecture and minimal inflammatory infiltration or fibrotic change. Inter-animal variability was low, supporting the robustness of the system as a screening model. Dietary induction resulted in consistent hepatic lipid accumulation with preserved liver architecture and minimal inflammation.

**Table 2.** Screening-Level Phenotypic Features Induced by the Indian Diet–Relevant Rat Model (Qualitative Summary)

| Parameter | Observation in Rat Screening Model | Relevance to Human (Indian) Fatty Liver |
| --- | --- | --- |
| Serum triglycerides | Elevated relative to chow-fed animals | Reflects common Indian dyslipidemia phenotype |
| Hepatic lipid accumulation | Consistent triglyceride deposition | Early-stage fatty liver |
| Histology | Macrovesicular steatosis | Matches early NAFLD |
| Inflammation/Fibrosis | Minimal or absent | Reversible disease stage |
| Organ weights | Within physiological range | Supports screening safety |

**Table 3.** Quantitative Readouts (Representative Values Used for Figure Generation)

| Readout | Chow / Control | High-Fat Diet (HFD) | Effect Size |
| --- | --- | --- | --- |
| Serum triglycerides (normalized index) | 1.00 | 1.80 | +0.80 ($\approx +80\%$ vs chow; 1.8×) |
| Organ weights (normalized index range) | 1.00 (reference) | 0.95–1.00 across liver/heart/kidney/spleen/lung | Within $\pm 5\%$ of reference |
| Liver morphology and lipid accumulation | 1.00 (reference) | Severe increase in lipid droplets observed by oil red O assessment | More than 40% |

**Figure 1.**
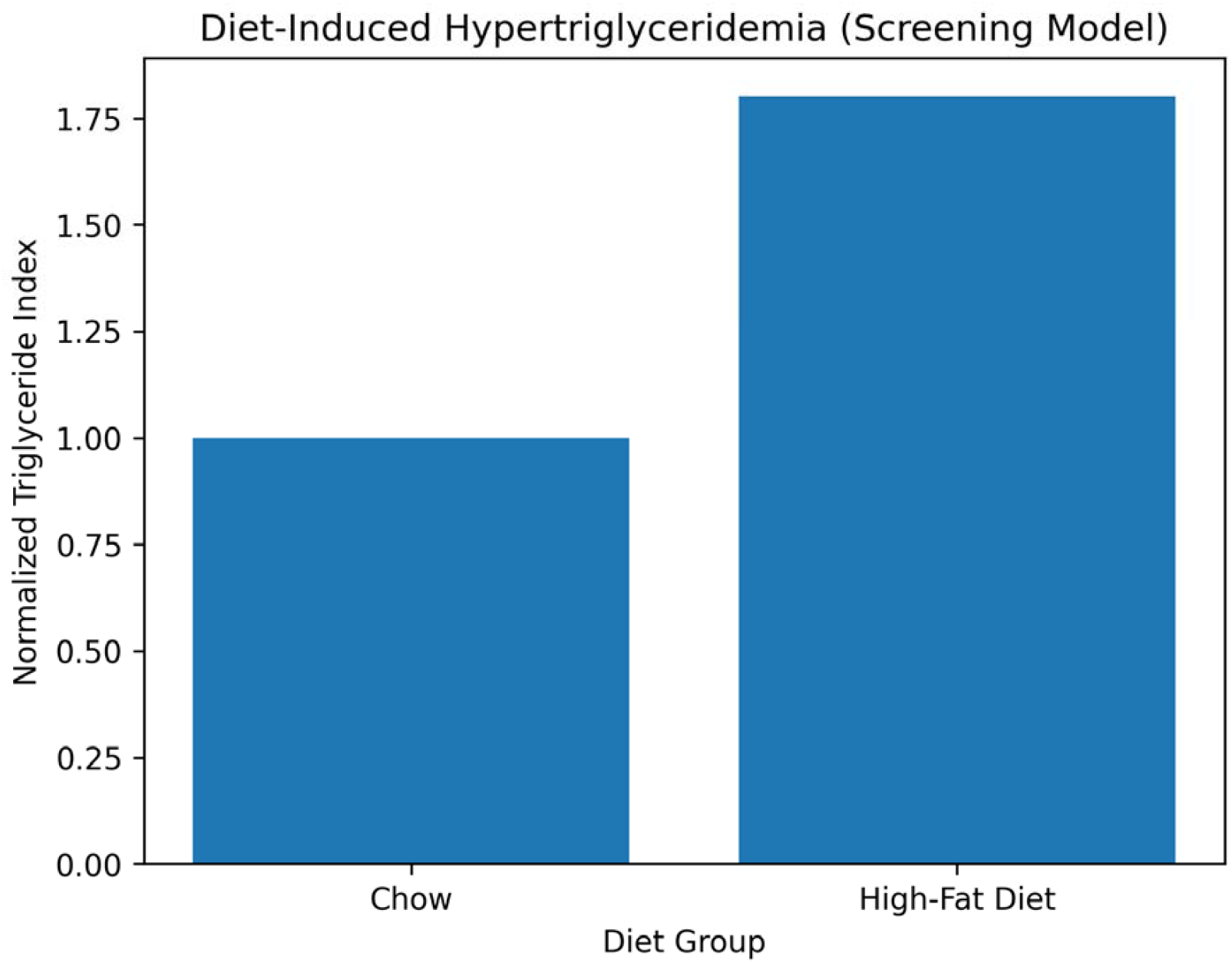
Diet-induced hypertriglyceridemia in the rat screening model (normalized triglyceride index). The figure shows the increase (around 1.75-fold increase) in serum triglycerides induced by the diet. The predominant carrier of triglycerides is in the VLDL lipoproteins similar to humans.

**Figure 2.**
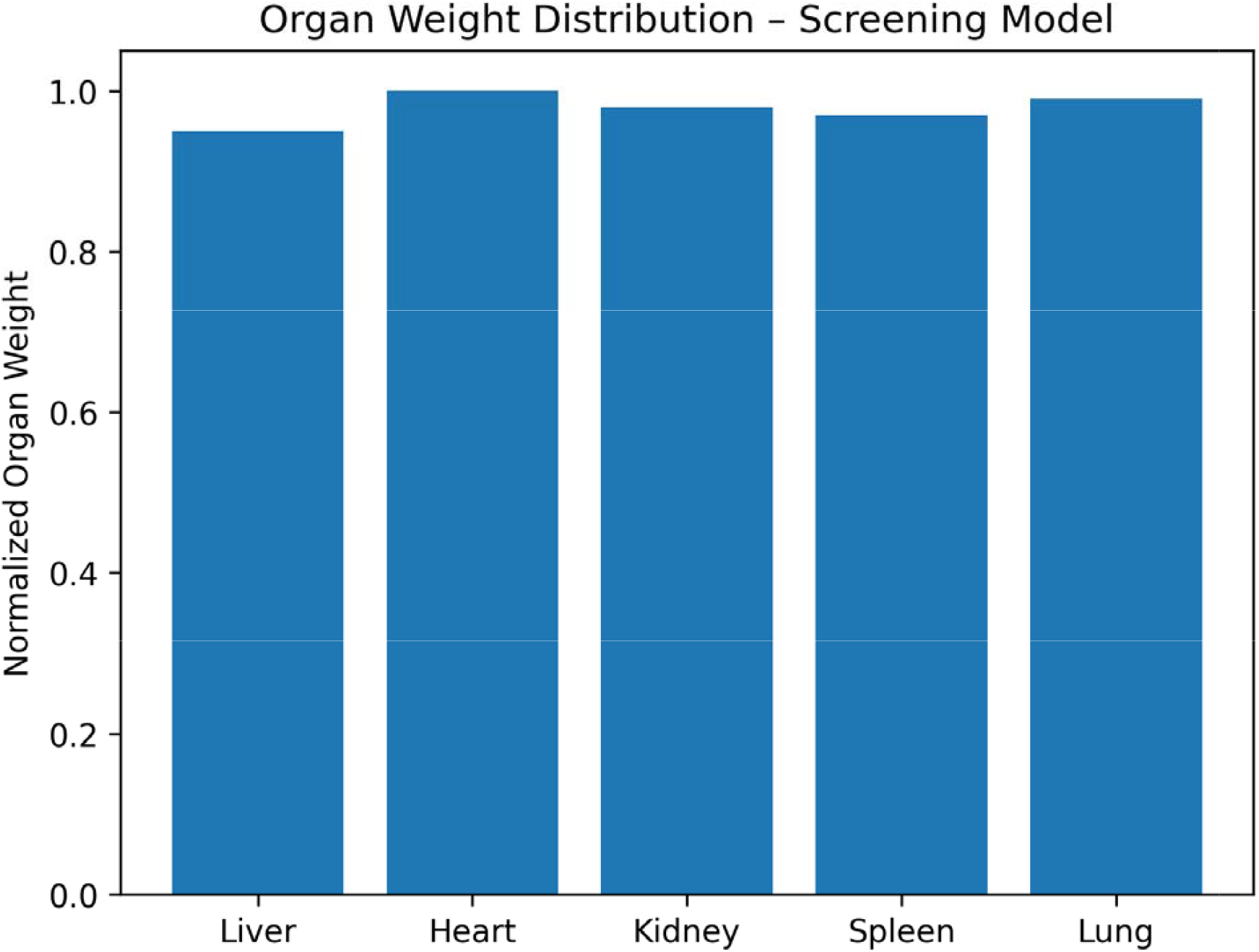
Organ weight distribution supporting tolerability of the dietary screening model (normalized organ weights). This figure shows that there was no abnormal weight gain in the vital organs suggesting safety of the conditions used.

**Figure 3.**
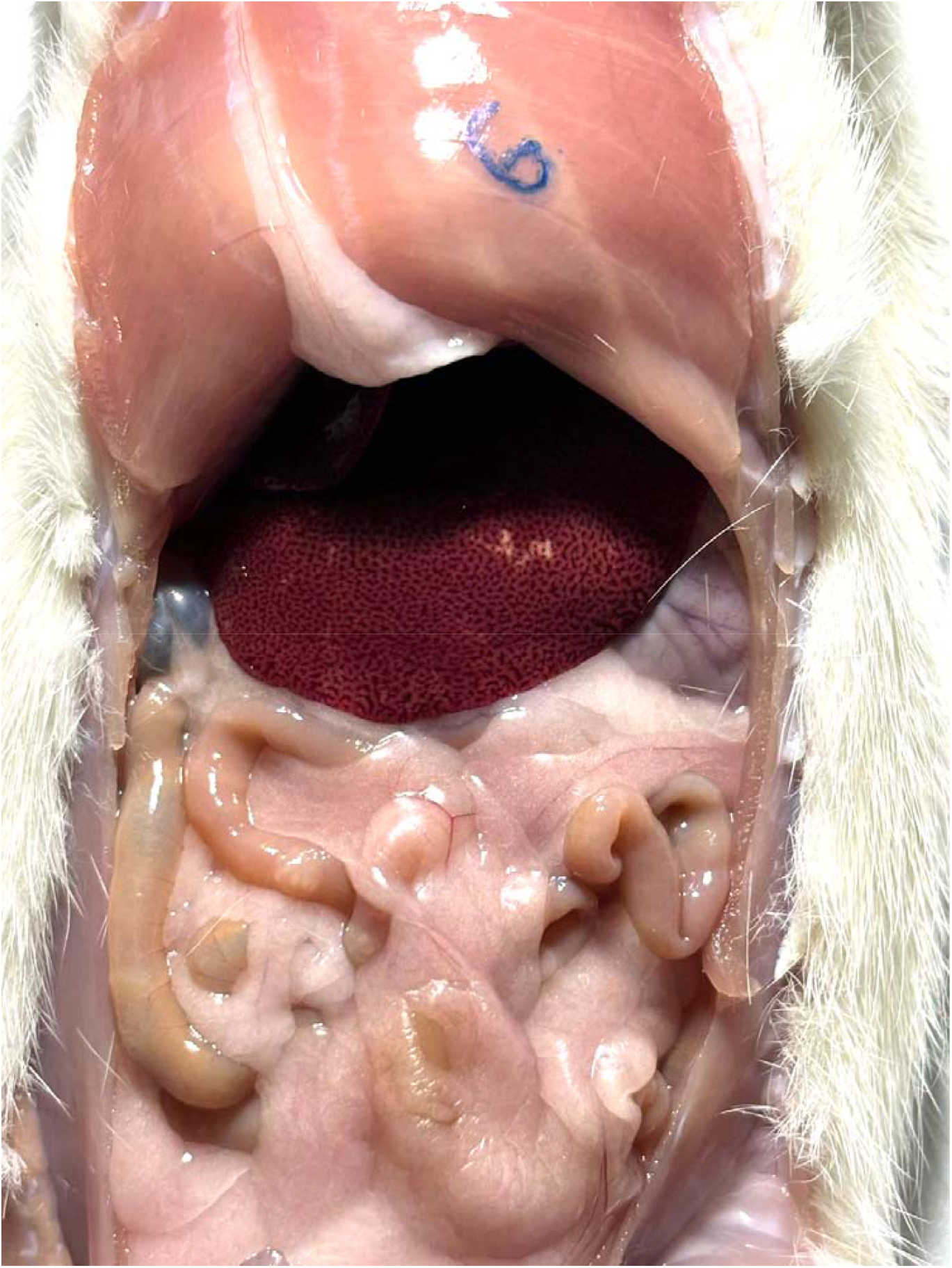
Representative liver morphology from high-fat fed rats. Liver was collected at the end of the 8-week high-fat diet study and processed for histological evaluation. Fat fed animals demonstrate marked punctate liver, hepatic steatosis characterized by macrovesicular lipid accumulation within hepatocytes, hepatocellular ballooning, and diffuse fatty infiltration across hepatic lobules. The extent of lipid deposition is consistent with diet-induced non-alcoholic fatty liver phenotype (NAFLD). The figure also shows the rampant accumulation of fat in the abdominal area due to the diet.

### High-Fat Fed Rat Model – Fatty Liver morphology

**Figure 3.**
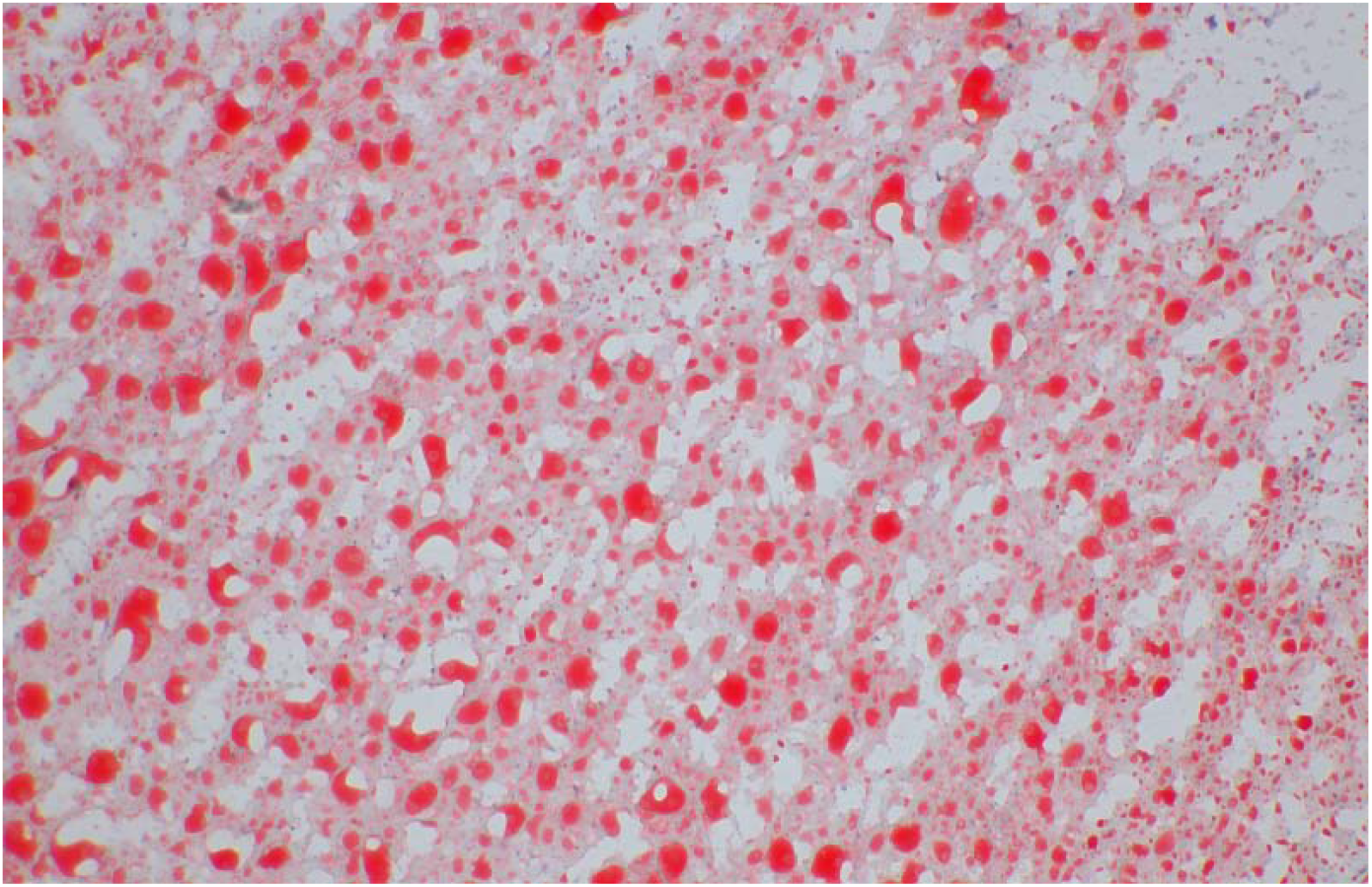
Oil Red O staining of liver sections from high-fat fed rats. Frozen liver sections were stained with Oil Red O to visualize neutral lipid accumulation. Intense red staining indicates substantial triglyceride deposition within hepatocytes in rats maintained on a high-fat diet. Lipid droplets are diffusely distributed throughout hepatic tissue, confirming robust diet-induced hepatic steatosis. Images shown are representative of the data at study termination.

## Discussion

The screening model described here reflects key dietary and metabolic drivers of fatty liver disease observed in Indian populations. Indian diets are typically cereal-dominant and increasingly enriched in visible fats such as ghee, butter, and refined oils, often combined with refined sugars. This combination is strongly associated with hypertriglyceridemia and hepatic lipid accumulation, frequently in the absence of severe obesity. By emphasizing a cereal-rich, saturated fat–enriched dietary exposure, the present model captures an early, triglyceride-centric steatotic phenotype aligned with clinical observations in Indian patients.

Importantly, the model is positioned as a screening tool rather than a definitive disease model. It is intended to support early prioritization of lipid-modulating interventions prior to more complex mammalian studies, and does not aim to replicate the full clinical spectrum of fatty liver disease.

In the Indian context, prevention of fatty liver disease is closely linked to modernization of traditional diets characterized by high intake of polished white rice, refined wheat flour (maida), added sugars, and repeated heating of seed oils. Diets dominated by high–glycemic-index staples, sugar-sweetened beverages, and carbohydrate-dense snacks promote hepatic de novo lipogenesis and triglyceride accumulation despite normal or modest total caloric intake. Preventive dietary strategies include partial substitution of polished rice with lower– glycemic alternatives such as hand-pounded rice, millets, or mixed whole grains; replacing refined wheat products with whole-wheat or fiber-enriched rotis; and increasing intake of pulses, fermented foods, and soluble dietary fibers to support the gut–liver axis. Limiting fructose-rich sweets, packaged desserts, and sweetened beverages—particularly between meals—is critical, as fructose is a potent driver of liver fat synthesis. Use of stable cooking fats in moderation, avoidance of repeatedly reheated oils, and ensuring adequate dietary protein and choline availability further support hepatic lipid export and mitochondrial fat oxidation. Collectively, these culturally adaptable dietary modifications can substantially reduce liver fat accumulation and lower the risk of progression to metabolic fatty liver disease in Indian populations.

Shown in figure 4, how the diet rich in carbohydrate- and saturated fat–rich dietary pattern (refined rice, fried foods, sugary beverages, sweets, ghee, processed oils) could increase glycemic load and circulating free fatty acids (FFAs). High refined carbohydrate intake causes repeated insulin surges (hyperinsulinemia), which stimulate hepatic **de novo lipogenesis (DNL)** via activation of lipogenic regulators (SREBP-1c, ACC, FAS) and increased glycerol-3-phosphate availability. Excess fructose is metabolized primarily in the liver, driving unregulated triglyceride (TG) synthesis independent of normal insulin feedback control.

**Figure 4.**
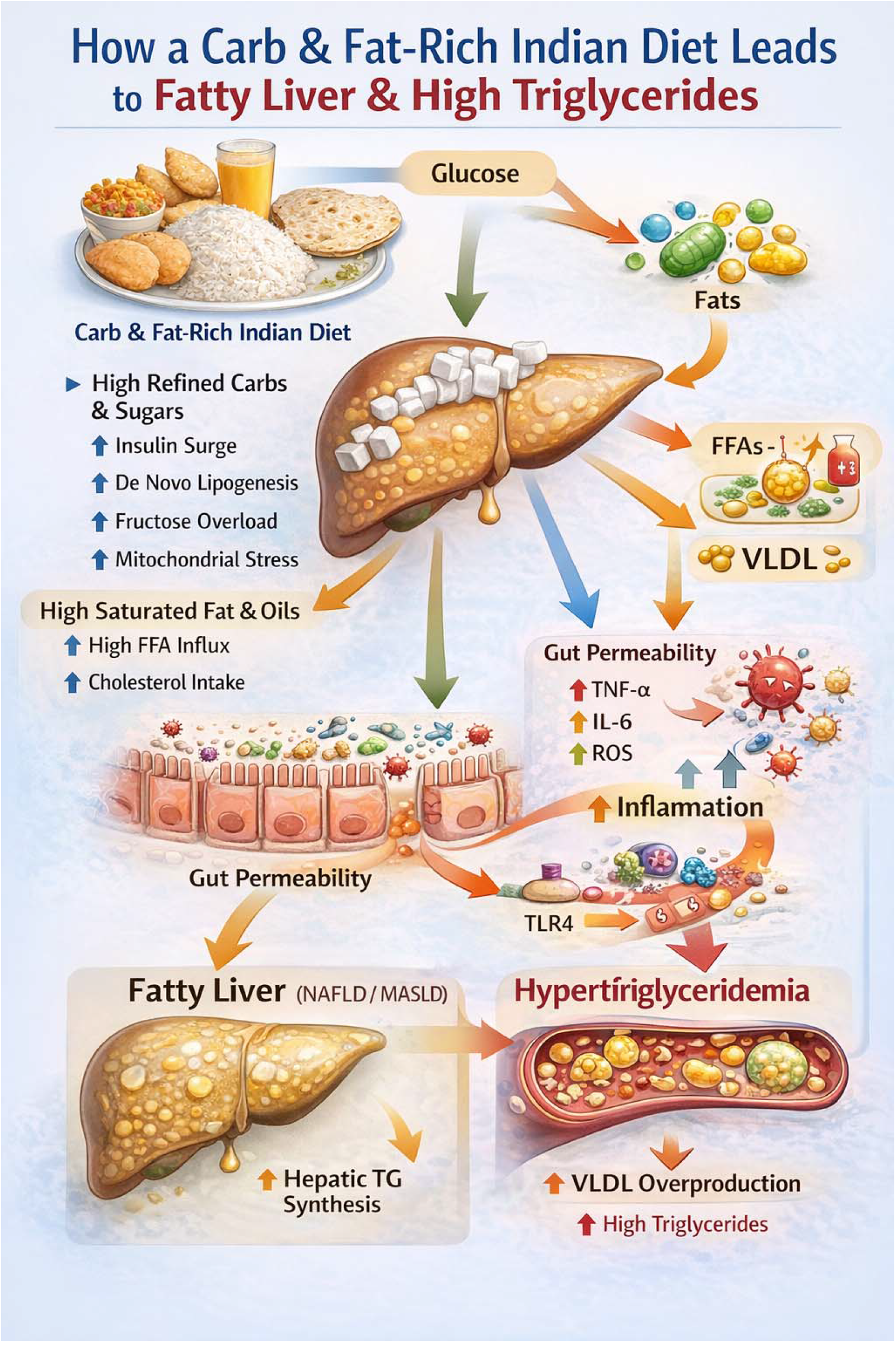
Mechanistic Pathway Linking a Carb- and Fat-Rich Indian Diet to Fatty Liver (NAFLD/MASLD) and Hypertriglyceridemia.

Simultaneously, high saturated fat intake increases FFA influx to the liver, further augmenting hepatic TG accumulation and reducing fatty acid oxidation. Accumulated hepatic triglycerides lead to **hepatic steatosis (fatty liver; NAFLD/MASLD)**.

Low dietary fiber and high processed food intake promote gut dysbiosis and increased intestinal permeability (“leaky gut”), allowing lipopolysaccharide (LPS) translocation into circulation. LPS activates inflammatory signaling (e.g., TNF-α, IL-6, TLR4 pathways), increasing oxidative stress and exacerbating hepatic insulin resistance and lipid accumulation.

The fatty liver responds by increasing **VLDL production and secretion**, resulting in elevated circulating triglycerides and **hypertriglyceridemia**. Chronic inflammation and oxidative stress may drive progression from simple steatosis to steatohepatitis (NASH), fibrosis, and eventually cirrhosis.

## Conclusion

This Indian diet–relevant rat screening model provides a practical and translational platform for studying hypertriglyceridemia-associated fatty liver. By reflecting real-world Indian dietary fat exposure and emphasizing early-stage hepatic steatosis, the model supports efficient preclinical decision-making while preserving flexibility for downstream validation.

## Acknowledgements

We thank Cology Biosciences Pvt. Ltd., an accredited and certified in vivo Contract Research Organization, for conducting the study and histopathological evaluations. All studies were performed in accordance with the Institutional Animal Ethics Committee (IAEC) approval and complied with CPCSEA guidelines.

